# Listening to bacterial Esperanto: transcriptome reprogramming in a plant beneficial rhizobacterium

**DOI:** 10.1101/2021.03.30.437723

**Authors:** Ana Bejarano, Michele Perazzolli, Ilaria Pertot, Gerardo Puopolo

## Abstract

Intraspecies, interspecies and interkingdom signalling occurring via diffusible communication signals (DCSs) continuously shapes the gene expression patterns of individual bacterial species in the rhizosphere, affecting bacterial functions within the rhizosphere microbial community. To unravel how DCSs influence rhizosphere competence of plant beneficial rhizobacteria, we carried out a functional and transcriptome analysis on the plant beneficial bacterium *Lysobacter capsici* AZ78 (AZ78). Results reveal that 13-methyltetradecanoic acid and indole, glyoxylic acid and 2,3-butanedione play a relevant role in the interaction between *L. capsici* members and other soil-living (micro)organisms. DCSs regulated mechanisms of multistress and multidrug tolerance and persistence, including detoxification, motility, antibiotic production, and expression of secretion systems. In particular, 13-methyltetradecanoic acid, glyoxylic acid and 2,3-butanedione might enable AZ78 to rapidly colonize the rhizosphere. Moreover, glyoxylic acid and 2,3-butanedione elicit biological responses to outcompete other (micro)organisms. In contrast, indole inactivates twitching motility and antibiotic production. These results demonstrate that DCSs influence the functioning of plant beneficial rhizobacteria and suggest that plant beneficial *L. capsici* strains could use them to foster rhizosphere colonization and enhance *in vivo* activities to increase soil health and plant fitness.

## Introduction

Rhizosphere microbial communities play a key role in soil fertility and in controlling soil-borne phytopathogenic microorganisms [1]. Their composition and functioning relay on root exudates and signalling molecules governing the complex interactions occurring between microorganisms and plants [2]. Bacterial intraspecies and interspecies communication occur mainly via Diffusible Communication Signals (DCSs) that allow bacterial communities to form and synchronize their behaviour [2]. N-acyl homoserine lactones (AHLs), produced by Gram-negative plant-associated bacteria, are among the most studied DCSs [3, 4]. Several works showed that endogenous and exogenous AHLs play an essential role in multiple bacterial physiological and biochemical behaviour [5]. Diffusible Signal Factors (DSFs) is another subgroup of DCSs produced by Gram-negative bacteria and it was initially reported in the phytopathogenic bacterium *Xanthomonas campestris* pv. *campestris* (*Xcc*) [6]. DSFs have been identified in other bacterial species as well and they have been linked to virulence, motility, biofilm production, and extracellular enzyme production [7]. Besides DSFs, Xanthomonadaceae family use diffusible factors (DFs) as DCSs; these signals are involved in the regulation of secondary metabolites biosynthesis and antioxidant activity [8–11]. In addition, plant-associated bacteria produce volatile organic compounds (VOCs) that are involved in communication and competition between physically separated soil microorganisms [12]. Among VOCs, indole (IND) is an ubiquitous interkingdom signal that influences antibiotic resistance, motility, biofilm formation, and virulence, and has the potential to be a DCS [13]. Other VOCs mediating changes in gene expression related to motility and antibiotic resistance are 2,3-butanedione (BUT) and glyoxylic acid (GLY) [14].

Among plant-associated Gram-negative bacteria, *Lysobacter* spp. belonging to the Xanthomonadaceae family, are commonly found in the plant rhizosphere where they control phytopathogenic microorganisms [15]. This ability mainly relies on the release of lytic enzymes and antibiotics like the Heat Stable Antifungal Factor (HSAF), which are toxic to phytopathogenic (micro)organisms [16, 17]. In *L. enzymogenes* DSFs, DFs and IND regulate HSAF biosynthesis [9, 18, 19] and twitching motility [20]. Furthermore, IND reverses the intrinsic antibiotic resistance of *L. enzymogenes* through the two-component regulatory system QseC/QseB [19, 20]. However, with the only exception of the involvement of DSFs and AHLs in *L. brunescens* behaviour [21], a complete overview of the overall effect of DCSs in *Lysobacter* spp. has not been described yet. In particular, it is not known if intra-, interspecies and interkingdom DCSs have a role in the establishment of *Lysobacter* spp. in the plant rhizosphere and if DCSs influence *Lysobacter* spp. to form stable communities with resident (micro)organisms or its ability to control phytopathogenic microorganisms.

To clarify these questions, we carried out a functional and transcriptome analysis of the plant beneficial bacterium *L. capsici* AZ78 (AZ78) in presence of a wide range of DCSs (13-methytetradecanoic acid [LeDSF3], IND, GLY, BUT, 3-hydroxybenzoic acid [3HBA], 4-hydroxybenzoic acid [4BHA], N-[3-hexanoyl]-L-homoserine lactone, N-[3-oxooctanoyl]-L-homoserine lactone, and N-[3-oxododecanoyl]-L-homoserine lactone). Genes encoding proteins involved in cell-cell communication systems were identified through genome mining. Functional experiments aimed to the assess changes in the biological properties of AZ78 upon exposure to DCSs and paid particular attention to bacterial growth and the ability of AZ78 to inhibit plant pathogens. Simultaneously, gene expression profiling of AZ78 exposed to main DCSs was carried out by high throughput RNA-Seq.

## Materials and methods

### Microorganisms and diffusible communication signals

Bacterial strains (Table S1) were routinely grown on Nutrient Agar (Oxoid, Basingstoke, UK) at 27°C. The phytopathogenic oomycete *Pythium ultimum* was maintained on Potato Dextrose Agar (Oxoid) at 25°C.

LeDSF3 was obtained from Avanti Polar Lipids (Alabaster, Alabama, USA). IND, GLY, BUT, 3HBA, 4BHA, N-(3-hexanoyl)-L-homoserine lactone, N-(3-oxooctanoyl)-L-homoserine lactone; and N-(3-oxododecanoyl)-L-homoserine lactone were purchased from Merck (Sigma-Aldrich, Darmstadt, Germany). Aqueous stock solutions were prepared, except for LeDSF3 and the mixture of AHLs that were prepared in pure methanol.

### Genome mining

AZ78 genome was mined to identify putative genes involved in cell-cell communication systems using nucleotide and protein sequence comparison. Genes from *L. enzymogenes* C3, *Stenotrophomonas maltophilia* (*Sm*) K279a and *Xcc* ATCC 33913^T^ were aligned against AZ78 genome, using RAST [22] to identify putative AZ78 genes responsible for DCS synthesis, reception and regulation using a cut-off of 1×10^−5^ at amino acid level. Putative genes were analysed with BLASTP [23], and length > 70 and identity > 70% at amino acid level were used as threshold. Identified gene clusters encoding putative proteins involved in cell-cell communication systems in AZ78 were then used to mine the *Lysobacter* spp. genomes, following the methodology described above. All genomes were downloaded from the National Center for Biotechnology Information (NCBI) (https://www.ncbi.nlm.nih.gov/) (Table S1). For the phylogenetic analyses, nucleotide sequences were aligned using ClustalW [24]. Evolutionary distances were assessed by applying Kimura’s two-parameter model [25] and the best phylogenetic trees were inferred by neighbour-joining method [26] implemented in MEGA 7 [27]. Confidence values for nodes in the trees were generated by bootstrap analysis [28] using 1000 permutations of the data sets.

### Assessment of diffusible communication signal production

Production of AHLs by AZ78 and *Lysobacter* spp. type strains was assessed by evaluating their ability to restore violacein production in *Chromobacterium violaceum* CV026 and/or to promote *lacZ* transcription in *Agrobacterium tumefaciens* NT1 (pZLR4) as previously described [29, 30]. Likewise, the ability to release DSF was determined using the bacterial reporter strain *Xcc* 8523 pL6engGUS according to Slater et al., 2000 [31]. *Pseudomonas chlororaphis* M71 [32] was used as an AHL positive control, whereas *Xcc* 8004 was used as a DSF positive control [6]. For each condition, five replicates were used, and the experiment was repeated.

### Evaluation of diffusible communication signal effect on cell growth

The effect of DCSs on AZ78 cell growth rate was assessed on 1/10 Tryptic Soy Broth (Oxoid) amended with each DCS (Table S2). AZ78 (starting concentration 1×10^7^ CFU/mL) was grown at 27°C on a 96-well plate (200 μL) and absorbance at 600 mm was recorded on a microplate reader (Synergy 2 Multi-Mode Microplate Reader, BioTek, Winooski, Vermont, USA). Non-inoculated media were used as blank. For each condition, five replicates were used. The experiment was repeated.

### Effect of diffusible communication signals on antimicrobial activity

The effect of DCSs on AZ78 antimicrobial activity was evaluated on Rhizosphere Mimicking Agar (RMA) [33] and the experimental design was made up to 8 treatments (Table S2). Inhibitory activity of AZ78 against *P. ultimum* was evaluated by using the classic dual-culture method. In brief, 10 μL of AZ78 cell suspension (1×10^8^ CFU/mL) were spot-inoculated at 3 cm of the edge of a plate. After 48 h incubation at 27°C, mycelium plugs (4 mm) were cut from the edge of one-week old *P. ultimum* plate, placed at 2.5 cm distance from AZ78 and incubated at 25°C for 168 h. AZ78 activity against *Rhodococcus fascians* LMG 3605 was determined by spot-inoculating 10 μl of AZ78 cell suspension (1×10^8^ CFU/mL) in the centre of a RMA plate. After 48 h incubation at 27°C, AZ78 cells were killed by exposure to chloroform vapour for 60 min [34]. Dishes were aerated under a laminar flow for 60 min, overlaid with 4 mL of 0.45% agar PBS containing *R. fascians* LMG 3605 (1×10^7^ CFU/mL) and incubated at 27°C for 72 h. RMA dishes seeded only with *P. ultimum* or *R. fascians* LMG 3605 were used as control.

Pictures were obtained with Bio-Rad Quantity One software implemented in a Bio-Rad GelDoc Imaging system (Bio-Rad Laboratories, Hercules, California, USA). Inhibitory activity was quantified by scoring *P. ultimum* or *R. fascians* LMG 3605 growth area (cm^2^) using ImageJ 1.52a [35] and calculated according to the formulas below:

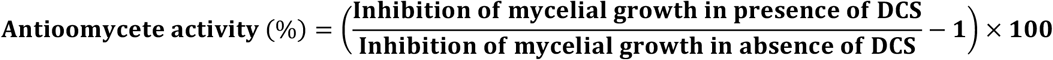

where

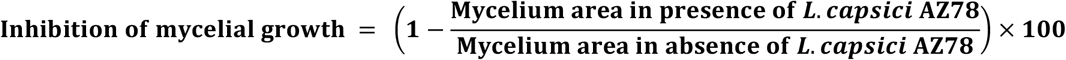

The effect of DCSs on AZ78 antibacterial activity was assessed as follows:

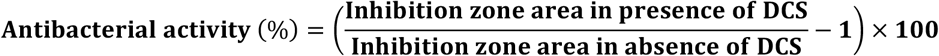

In both cases, treatments included five replicates and experiments were repeated.

### RNA extraction

The AZ78 response to DCSs was evaluated on RMA and the experimental design was made up to 8 treatments in triplicate (Table S2). Ten microliters of AZ78 cell suspension (1×10^10^ CFU/mL) were spot-inoculated in the centre of a RMA plate and incubated at 27°C for 48 h. Plugs (7 mm diameter) were collected from the AZ78 macrocolonies, immediately frozen in liquid nitrogen and stored at −80°C. Frozen samples were processed according to Brescia et al., [33] and total RNA was extracted using Spectrum Plant Total RNA Kit (Sigma-Aldrich). DNase treatment was performed with the RNase-Free DNase set (Qiagen, Hilden, Germany). RNA integrity and concentration were assessed using a 2200 TapeStation System (Agilent Technologies, Santa Clara, California, USA) and a Qubit 4 Fluorometer (ThermoFisher Scientific, Carlsbad, California, USA) with Qubit RNA BR assay kit (ThermoFisher Scientific), respectively.

### Illumina sequencing and mapping to the reference genomes

Library construction and Illumina Sequencing were carried out at Fasteris (Plan-les-Ouates, Switzerland). Ribosomal RNA (rRNA) depletion was performed using the Ribo-Zero rRNA Removal Kits (Bacteria) (Illumina, San Diego, California, USA). Complementary DNA (cDNA) libraries were synthesised using TruSeq Stranded mRNA Library Prep (Illumina, USA), they were multiplexed (two libraries per lane) and paired-end reads of 150 nucleotides were obtained using an Illumina HiSeq 4000 (Illumina), resulting in ∼7-42 million reads per sample (Table S3). Raw sequences were deposited at the Sequence Read Archive of the NCBI under BioProject number PRJNA714393.

Sequence analysis was carried out using Omicsbox 1.3.11 (www.biobam.com/omicsbox). Illumina HiSeq data was assessed for quality using FastQC [36]. Raw reads for each sample were trimmed to increase overall quality using Trimmomatic 0.38 [37]. The resulting reads were aligned to AZ78 genome (Table S1) using the STAR 2.7.5a [38] and read counts were extracted from STAR alignments using HTSeq [39].

### Identification of differentially expressed genes and functional annotation of RNA-Seq

Genes with zero counts in all replicates were excluded from the analysis and raw counts were normalised using the trimmed mean of M-values method [40]. Differentially expressed genes (DEGs) were identified using edgeR 3.28.0 [41] using a p-value < 0.01 and a log fold change (FC) of at least 1-fold upregulation/downregulation as cut off values. Venn diagrams summarizing DEGs distribution were drawn with VennPainter [42]. Hierarchical clustering and heat maps were created with TreView3 [43].

The protein sequences of all predicted genes [34] were functionally annotated using Blast2Go (http://www.blast2go.org) [44]. Default settings were applied and a minimum E-value of 10^−5^ was imposed as cut off. DEGs were further annotated based on the NCBI gene description and classified in 20 functional categories.

### Validation of RNA-Seq

First-strand cDNA was synthetized from 600 ng of purified RNA with SuperScript III Reverse Transcriptase (Invitrogen, Carlsbad, California, USA) using random hexamers, according to manufacturer’s instructions. qRT-PCR reactions were carried out with Platinium SYBR Green qPCR Super-Mix-UDG (Invitrogen, USA) and specific primers (Table S4) were designed using Primer3 software [45]. Primer specificity was assessed using PCR before gene expression analysis. qRT-PCR reactions were run for 50 cycles (95 °C for 15 s and 60 °C for 45 s) on a LightCycler 480 (Roche Diagnostics, Mannheim, Germany). Each sample was examined in three technical replicates and dissociation curves were analysed to verify the specificity of each amplification reaction. LightCycler 480 software, version 1.5 (Roche Diagnostics, Mannheim, Germany) was used to extract cycle threshold (Ct) values based on the second derivative calculation and the LinReg software, version 11.0, was used to calculate reaction efficiencies for each primer pair [46]. Relative expression levels were calculated according to Pfaffl equation [47] using AZ78 growing in RMA as calibrator. The housekeeping gene *recA* (AZ78_1089; [34]) was used as constitutive gene for normalization. The linear relationship between the RNA-Seq log_2_FC values and the qRT-PCR log_2_FC values of selected genes was estimated by Pearson correlation analysis.

### Statistical analysis

Percentage values were arcsine square root transformed to normalize distributions and to equalize variances. Comparisons between repeated experiments of antimicrobial activity were done using two-way analysis of variance (ANOVA) and the data were pooled when no significant differences were found according to the *F*-test (*p* > 0.05). Data were analysed using one-way ANOVA and Tukey’s test (α = 0.05) was used to detect significant differences. Statistical analyses were carried out using IBM SPSS Statistics for Windows, Version 21.0 (IBM Corp, Armonk, New York, USA).

## Results

### Cell-cell communication systems in *Lysobacter capsici* AZ78 genome

Putative genes encoding the DSF response regulator RpfG (AZ78_0630) and the turn-over signal RpfB (AZ78_0629) were found in the AZ78 genome and showed an identity greater than 80% (at amino acid level) with its homologues in *L. enzymogenes* C3*, Sm* K279a, and *Xcc* ATCC 33913^T^ (Table 1). *rpfG* and *rpfB* genes in AZ78 were closely related to their homologues in other *Lysobacter* spp. and clustered together (Figure S1). AZ78 genome also included a putative DSF sensor RpfC (AZ78_3298) and a crotonase/enoyl-Coenzyme A (RpfF) (AZ78_3297) with low similarity compared to *L. enzymogenes* C3*, Sm* K279a, and *Xcc* ATCC 33913^T^ (Table 1, Figure S2). The *rpfF/rpfC* region (3,947,548 – 3,948,450 bp) was located far from the *rpfG/rpfB* region (857,114 – 859,152 bp) in AZ78 (Figure 1a). DSF biosynthesis was confirmed by AZ78 ability to induce the glucuronidase activity in *Xcc* 8523 pL6engGUS like the control strain *Xcc* 8004 (Figure 1b).

**Table 1.**
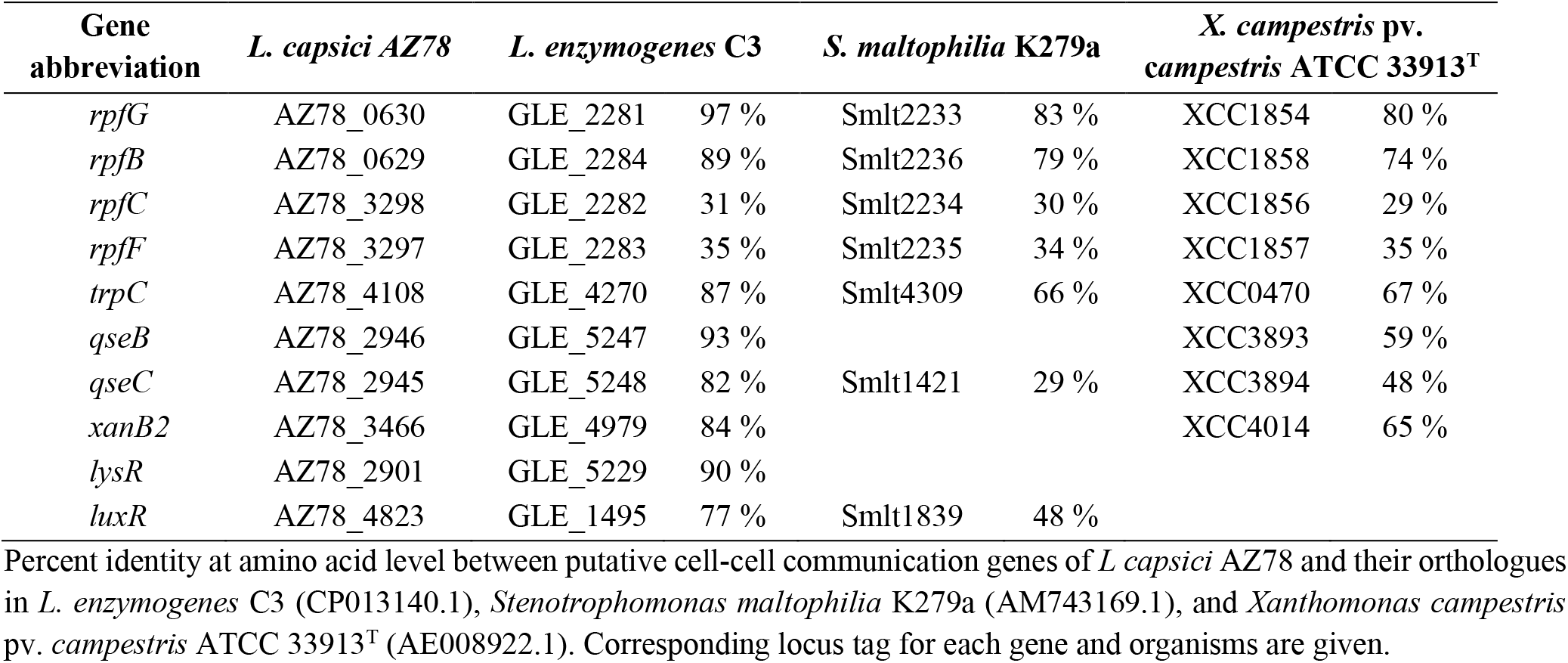
Cell-cell communication genes present in the genome of *Lysobacter capsici* AZ78.

**Figure 1.**
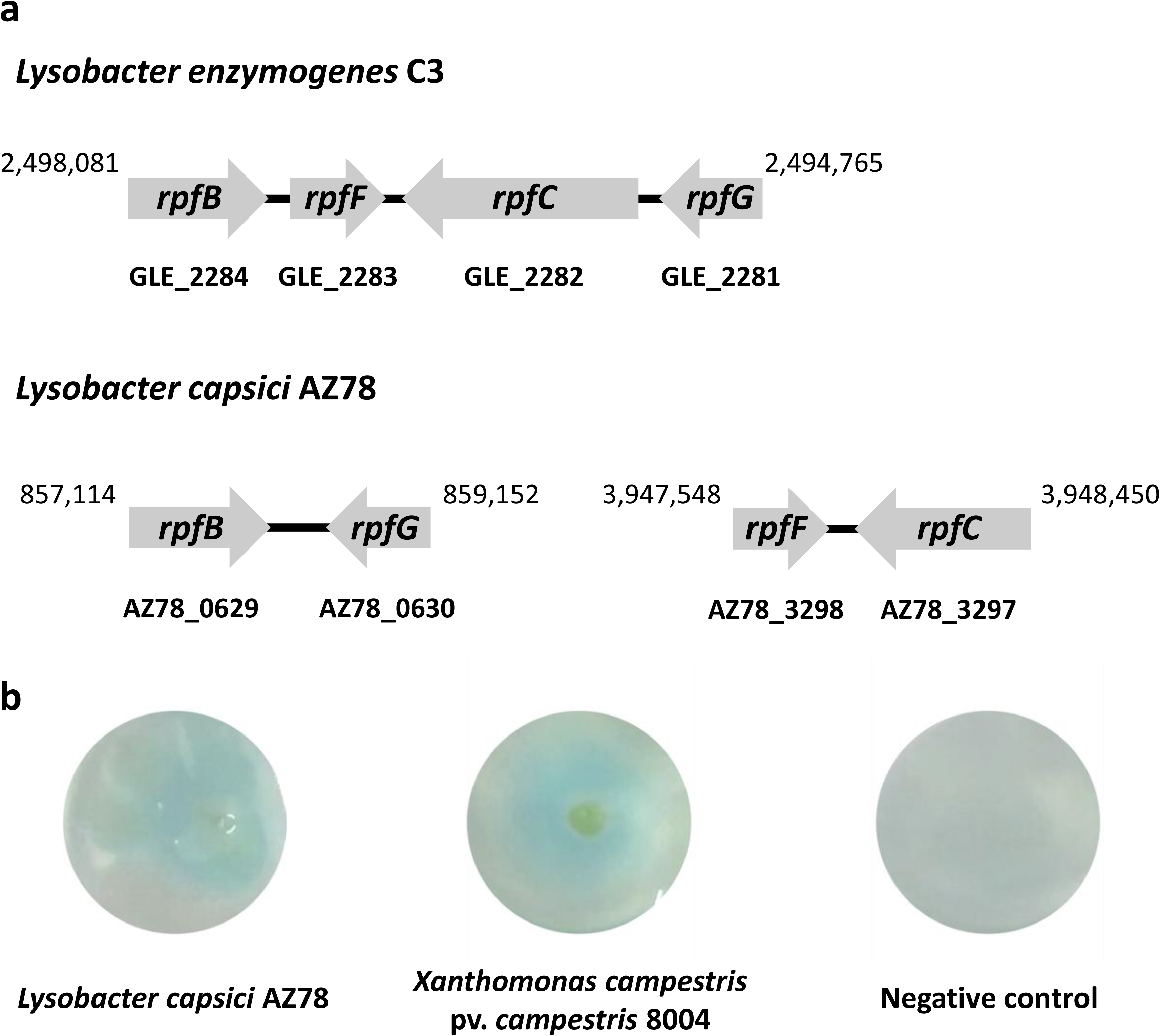
Location of *rpf* genes in *Lysobacter capsici* AZ78. (a) Comparison of the clusters of genes involved in Diffusible Soluble Factors synthesis and perception in *L. enzymogenes* C3 and *L. capsici* AZ78. Gene names are given within the arrows. The corresponding accession number is given under each gene. (b) Bioassay showing DSF production ability (blue halo) of *L. capsici* AZ78 in comparison to *Xanthomonas campestris* pv. *campestris* 8004.

As for VOCs, a putative gene encoding IND synthase (AZ78_4108) was found in AZ78 genome. Homologues of *qseB* (AZ78_2946) and *qseC* (AZ78_2945) involved in IND regulation were also found in AZ78 genome and showed high identity with *L. enymogenes* C3 homologue (Table 1). Homologues of IND synthase and QseB/QseC system were also present in other *Lysobacter* spp. (Figure S3).

A gene encoding a putative chorismatase needed for DFs production was found in AZ78 genome (AZ78_3466) with an identity of 64% at amino acid level with the homologue *xanB2* (XCC4014) of *Xcc* ATCC 33913^T^. Furthermore, a homologue of the LysR family transcription factor involved in the DF regulatory cascade, was identified in AZ78 genome (AZ78_2901) (Table 1). Chorismatase and LysR transcription factor were also typical among other *Lysobacter* spp. (Figure S4).

AZ78 *luxR* gene (AZ78_4823), responsible for the detection and response to AHLs, had an amino acid identity of 77 and 48 with the homologues of *L. enzymogenes* C3 and *Sm* K279a, respectively (Table 1). *luxR* phylogenetic analysis showed that AZ78 *luxR* homolog clustered together with other members of *L. capsici* (Figure S4). LuxR was not associated to its cognate AHL synthase (LuxI) in AZ78 or in other *Lysobacter* species, with only exception of *L. daejeonensis* GH1-9^T^ having a *luxI* homologue. The absence of LuxI homologs was confirmed by the inability of AZ78 to restore β-galactosidase activity and violacein production in the reporter strains *A. tumefaciens* NT1 pZRL4 and *C. violaceum* CV026, respectively (Figure S5). In contrast, *L. daejeonensis* GH1-9 ^T^ was able to restore violacein production in *C. violaceum* CV026 confirming relation between the presence of *luxI* and AHL production.

### Diffusible communication signals affect *Lysobacter capsici* AZ78 growth and antimicrobial activity

GLY, BUT, 3HBA, 4HBA and AHL had no effect on AZ78 growth curves, whereas LeDSF3 and IND slowed down AZ78 growth compared to the untreated control (Figure S6).

Exogenous addition of DCSs to RMA showed no effect on *P. ultimum* and *R. fascians* LMG 3605 growth, but it modulated AZ78 antimicrobial activity against *P. ultimum* and *R. fascians* LMG 3605 (Figure 2). LeDSF3, 4HBA, and IND decreased AZ78 inhibitory activity against *P. ultimum* by a 5, 22 and 47 %, respectively. In contrast GLY, BUT, 3HBA, and AHL increased AZ78 inhibitory activity against *P. ultimum* up to a 7% (Figure 2). LeDSF3, IND, 3HBA, 4HBA and AHL decreased AZ78 inhibitory activity against *R. fascians* LMG 3605 up to a 31%, while BUT and GLY increased AZ78 antibacterial activity by a 9 and 48% respectively (Figure 2). Changes were particularly relevant for IND and GLY.

**Figure 2.**
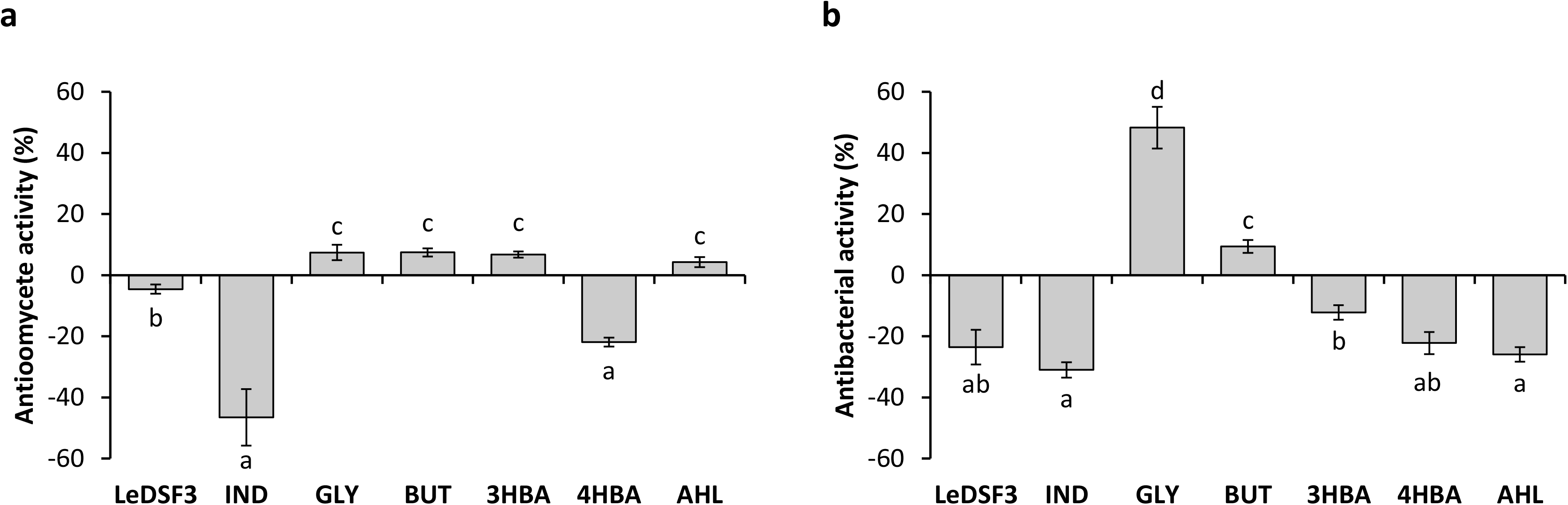
Effect of diffusible communication signals on the inhibitory activity of *Lysobacer capsici* AZ78 against (a) *Pythium ultimum* and (b) *Rhodococcus fascians*. Antioomycete and antibacterial activity is expressed as the mean value and standard error variation (percentage) of the reduction of the mycelium growth area of *P. ultimum* and *R. fascians* compared to the control (*L. capsici* AZ78 in not supplemented media), respectively. LeDSF3: 13-methyltetradecanoic acid 50 μM, IND: indole 500 μM, GLY: glyoxylic acid 0.01 μM, BUT: 2,3-butanedione 0.01 μM, 3HBA: 3-hydroxybenzoic acid 30 μM, 4HBA: 4-hydroxybenzoic acid 50 μM, AHL: mix of N-acyl homoserine lactones 20 μM. Each treatment included five replicates and data originating from two independent experiments were pooled. Different letters indicate significant letters according to Tukey’s test (α = 0.05).

### Transcriptional response of *Lysobacter capsici* AZ78 to diffusible communication signals

The expression of 21% of all AZ78 genes was significantly affected by DCSs (|log_2_FC| > 1 and *p* < 0.01). The largest number of DEGs, 636 genes (about 11.9% of AZ78 transcriptome), was found upon exposure of AZ78 to LeDSF3 (Table 2). This was followed by IND (603 DEGs, 11.3% of total genes), GLY (292 DEGs, 5.5% of total genes), BUT (237 DEGs, 4.4% of total genes), 3HBA (101 DEGs, 1.9% of total genes), and 4BHA (58 DEGs, 1.1% of total genes). The lowest number of DEGs (24, 0.5% of total genes) occurred among cells treated with AHL (Table 2). RNA-Seq results were validated by the relative expression level of 10 selected genes assessed using qRT-PCR (Table S4). A close correlation (Pearson’s *r* = 0.95) was observed between log_2_FC measured with RNA-Seq and qRT-PCR (Figure S7).

**Table 2.** Differentially expressed genes in Lysobacter capsici AZ78 in response to diffusible communication signals after 48 h incubation.

|  |  | LeDSF3 | IND | GLY | BUT | 3HBA | 4HBA | AHL |
| --- | --- | --- | --- | --- | --- | --- | --- | --- |
| UP-REGULATED | *Genes with assigned function* | 325 | 312 | 142 | 107 | 48 | 34 | 17 |
|  | *Hypothetical proteins* | 40 | 56 | 42 | 38 | 12 | 14 | 3 |
|  | **Total no. of genes** | **365** | **368** | **184** | **145** | **60** | **48** | **20** |
| DOWN-REGULATED | *Genes with assigned function* | 194 | 169 | 78 | 72 | 33 | 8 | 2 |
|  | *Hypothetical proteins* | 77 | 66 | 30 | 20 | 8 | 2 | 2 |
|  | **Total no. of genes** | **271** | **235** | **108** | **92** | **41** | **10** | **4** |
| TOTAL | *Genes with assigned function* | 519 | 481 | 220 | 179 | 81 | 42 | 19 |
|  | *Hypothetical proteins* | 117 | 122 | 72 | 58 | 20 | 16 | 5 |
|  | **Total no. of genes** | **636** | **603** | **292** | **237** | **101** | **58** | **24** |

Each treatment was subjected to pairwise comparison with the untreated control. |Log2-fold change| > 1 and p-value < 0.01 were chosen as cut-off values for differential gene expression. LeDSF3: 13-methyltetradecanoic acid 50 µM, IND: indole 500 µM, GLY: glyoxylic acid 0.01 µM, BUT: 2,3-butanedione 0.01 µM, 3HBA: 3-hydroxybenzoic acid 30 µM, 4HBA: 4-hydroxybenzoic acid 50 µM, AHL: mix of N-acyl homoserine lactones 20 µM.

Venn diagrams (Figure S8) revealed overlaps among the seven conditions, but it did not identified genes modulated by all seven DCSs. Moreover, a heatmap (Figure 3) showed that GLY and BUT clustered together, likewise 3HBA, 4HBA, and AHL. Instead, LeDSF3 and IND grouped independently.

**Figure 3.**
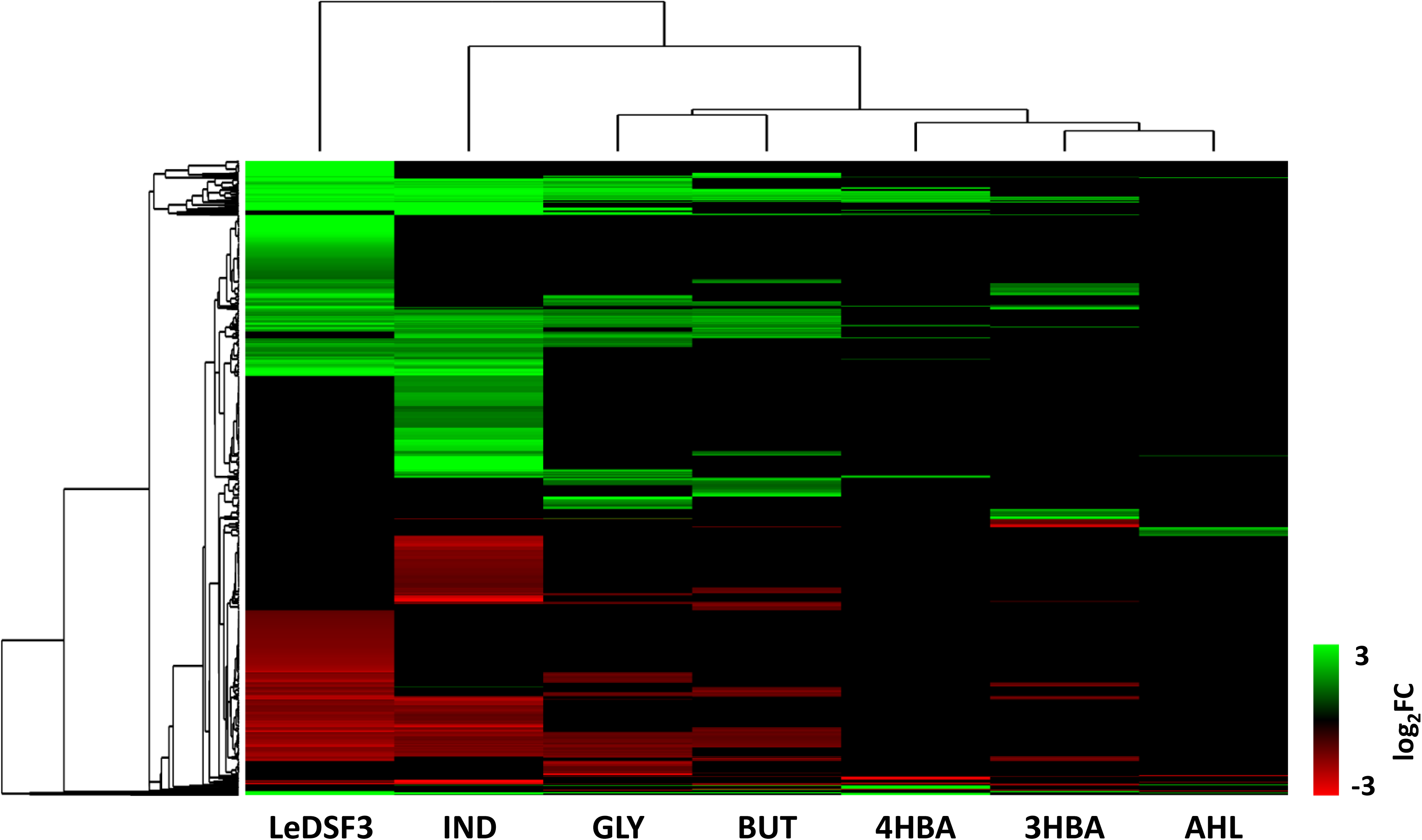
Hierarchical clustering of *Lysobacter capsici* AZ78 genes differentially expressed (1,115 genes, Tables S5-S11 for gene expression data). LeDSF3: 13-methyltetradecanoic acid 50 μM, IND: indole 500 μM, GLY: glyoxylic acid 0.01 μM, BUT: 2,3-butanedione 0.01 μM, 3HBA: 3-hydroxybenzoic acid 30 μM, 4HBA: 4-hydroxybenzoic acid 50 μM, AHL: mix of N-acyl homoserine lactones 20 μM.

### The active response of *Lysobacter capsici* AZ78 to diffusible communication signals

Functional annotation of AZ78 genes modulated by DCSs revealed that up-regulated DEGs were mainly related to global metabolism, growth, RNA transcription and degradation; transport, phosphotransferase systems, and secretion (Figure 4). Conversely, down-regulated DEGs were mainly related to DNA metabolism.

**Figure 4.**
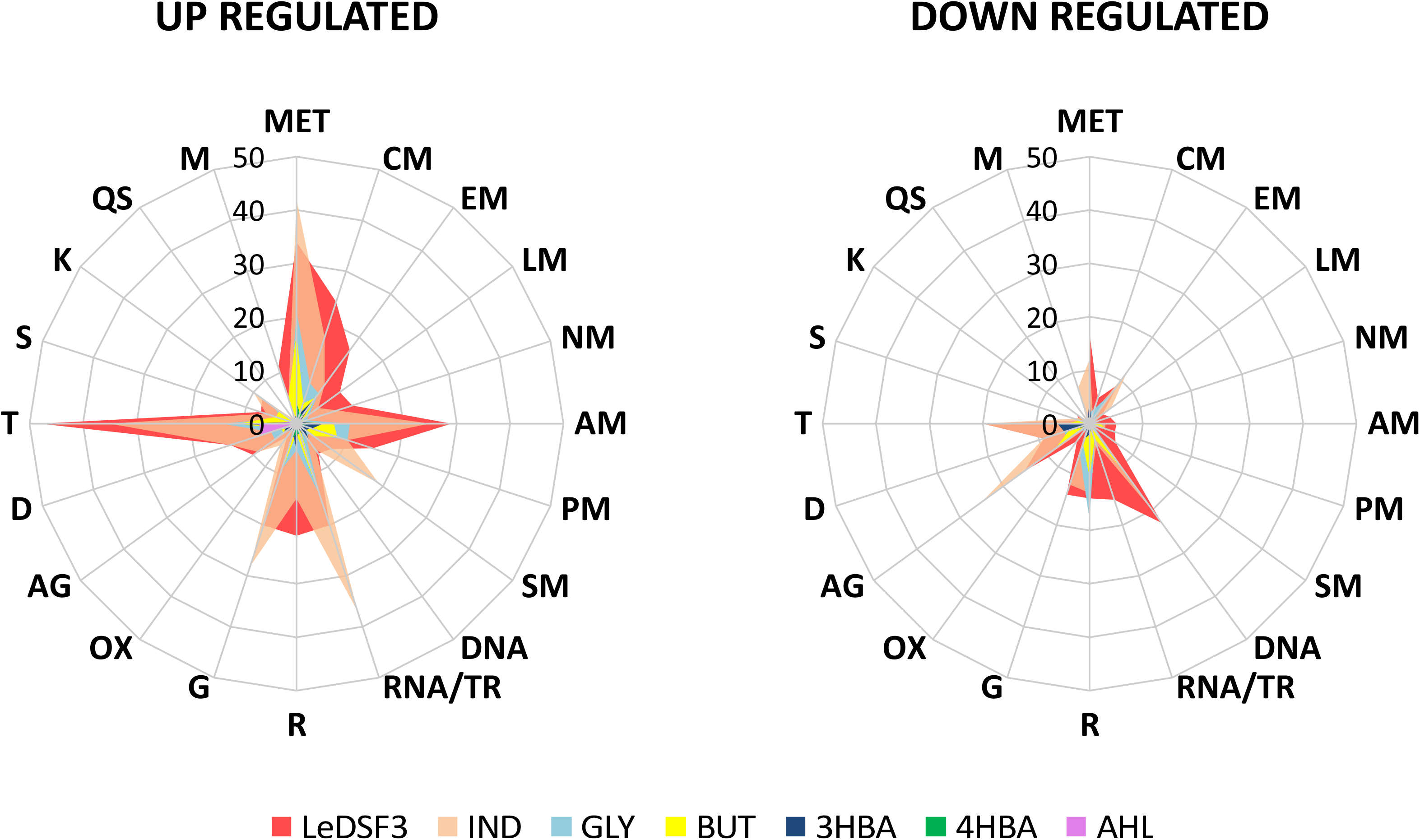
Radar plots of *Lysobacter capsici* AZ78 genes differentially expressed in response to diffusible communication signals. Annotated genes were classified in 20 functional categories: global metabolism (MET); carbohydrate metabolism (CM); energy metabolism (E); lipid metabolism (LM); nucleotide metabolism (NM); amino acid metabolism (AM); protein metabolism (PM); secondary metabolism (SM); DNA metabolism (DNA); RNA transcription and degradation (RNA/TR); translation (T); growth (G); oxidative stress (OX); antagonism (AG); defence (D); transport, phosphotransferase systems and secretion (T); signal transduction and receptors (S); kinase/phosphatase (K); quorum sensing (QS); motility, chemotaxis, and biofilm (M). Only genes with |log2-fold change| > 1 and p-value < 0.01 were included. LeDSF3: 13-methyltetradecanoic acid 50 μM, IND: indole 500 μM, GLY: glyoxylic acid 0.01 μM, BUT: 2,3-butanedione 0.01 μM, 3HBA: 3-hydroxybenzoic acid 30 μM, 4HBA: 4-hydroxybenzoic acid 50 μM, AHL: mix of N-acyl homoserine lactones 20 μM.

LeDSF3 regulated a total of 68 genes involved in transport, phosphotransferase systems, and secretion; 51 genes involved in global metabolism; 35 genes involved in RNA transcription and degradation; 35 genes involved in translation, 34 genes involved growth, and 34 genes involved in amino acid metabolism (Figure 4, Table S5). The category of transport, phosphotransferase systems, and secretion was affected by IND (55 genes) (Figure 4, Table S6), followed by global metabolism (54 genes), RNA transcription and degradation (41 genes), growth (40 genes), and antagonism (34 genes). GLY, BUT, and 3HBA modulated a relevant number of genes related to global metabolism and transport, phosphotransferase systems, and secretion (Figure 4, Tables S7, S8, S9). Moreover, GLY and BUT modulated a relevant number of genes classified into RNA transcription and degradation and translation, among which tRNA genes were mainly down-regulated. Genes related to transport, phosphotransferase systems, and secretion and defence were modulated by 4HBA and AHL (Figure 4, Tables S10, S11).

### Efflux pumps, type IV pilus biogenesis, antagonism, cell-cell communication systems, and secretion systems are influenced by diffusible communication signals in *Lysobacter capsici* AZ78

Many genes ascribed to transport, phosphotransferase systems, and secretion were involved in drug and metal (particularly iron) transport (Tables S5-S11). Major Facilitator Superfamily (MFS) transporters, like the multidrug exporter gene *yceL* (AZ78_2682), were up-regulated by LeDSF3 (Figure 5). Resistance-Nodulation-Division (RND) efflux system genes were up-regulated by DCSs, especially by AHL (Figure 5, Table S11). TonB-dependent receptors involved in the uptake of iron-siderophore complexes or vitamins were up-regulated by LeDSF3 and down-regulated by IND (Tables S5, S6). Additionally, DCSs up-regulated a relevant set of transcription regulators belonging to the AraC, ArsR, TetR, MerR and MarR families (Tables S5-S11).

**Figure 5.**
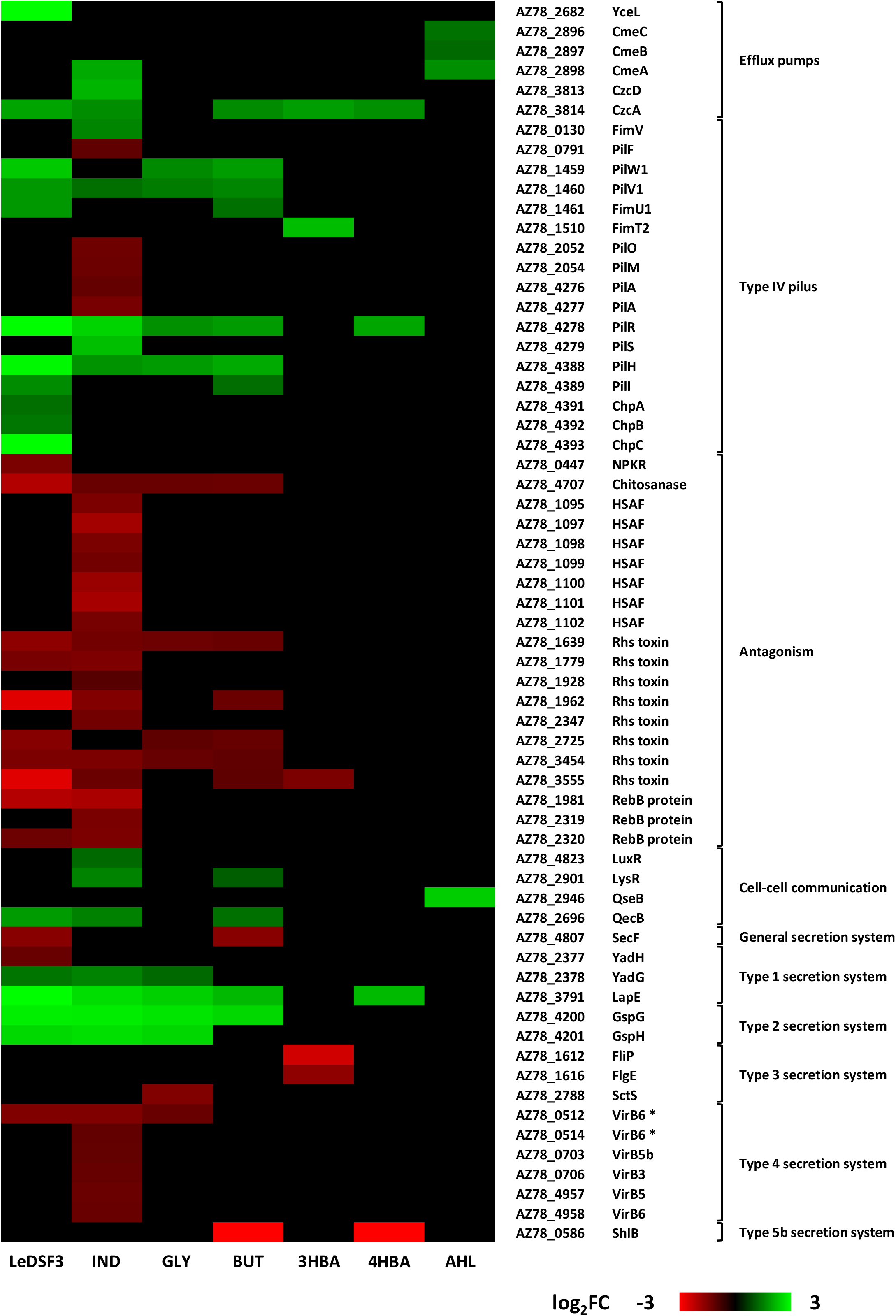
Heatmap of *Lysobacter capsici* AZ78 genes differentially expressed in response to diffusible communication signals and related to efflux pumps, type IV pilus, antagonism, cell-cell communication systems and secretion systems. HSAF: Heat Stable Antifungal Factor. VirB6 * represent duplicate gene pairs located through AZ78 genome. LeDSF3: 13-methyltetradecanoic acid 50 μM, IND: indole 500 μM, GLY: glyoxylic acid 0.01 μM, BUT: 2,3-butanedione 0.01 μM, 3HBA: 3-hydroxybenzoic acid 30 μM, 4HBA: 4-hydroxybenzoic acid 50 μM, AHL: mix of N-acyl homoserine lactones 20 μM.

With the only exception of AHL, DCSs modulated the expression of genes involved in type IV pilus (T4P) biosynthesis. LeDSF3 up-regulated the expression of *pilW1-pilV1-fimU1* genes (AZ78_1459-AZ78_1461), related to minor pilins; *pilR* from two-component system *pilR-pilS* (AZ78_4278-AZ78_4279); and *pilH* (AZ78_4388)*, pilI* (AZ78_4389)*, chpA* (AZ78_4391)*, chpB* (AZ78_4392) and *cphC* (AZ78_4393) from the pilus-specific chemotaxis system (Pil-Chp) (Figure 5, Table S5). Genes encoding minor pilins, pilRS, and Pil-Chp were also up-regulated in GLY, BUT, and 4HBA (Figure 5; Tables S7, S8, S10). IND up-regulated the expression of *fimV* (AZ78_0130), *pilV1*, *pilR-pilS*, and *pilH*, while it down-regulated *pilF* (AZ78_0791), *pilO* and *pilM* (AZ78_2052 and AZ78_2054), and the major pilin *pilA* (AZ78_4276-4277) (Figure 5, Table S6). LeDSF3 and IND down-regulated a relevant number of involved in antagonism. IND down-regulated the biosynthetic gene cluster AZ78_1095-AZ78_1102, responsible for the production of HSAF [33] (Figure 5; Table S6), and several extracellular proteases (e.g. AZ78_2802 and AZ78_3223) (Table S6). Moreover, IND up-regulated a diguanylate cyclase harbouring a GGDEF motif (AZ78_4062) possibly related to cyclic-di-GMP (c-di-GMP) biosynthesis (Table S6). Other genes related to antagonism, such as genes encoding Rhs toxins and RebB proteins, responsible for the expression of killing traits, were down-regulated by LeDSF3 and IND (Figure 5; Tables S5, S6).

DCS also caused changes in the expression of genes involved in the reception and regulation of DCSs in AZ78 (Figure 5). For instance, the transcription factor LysR (AZ78_2901) was up-regulated by IND and BUT, *qseB* (AZ78_2946) was up-regulated by AHL and *luxR* (AZ78_4823) by IND.

Genes related to Type I secretion system (T1SS) were mostly up-regulated by all DCSs, especially by LeDSF3 and IND (Figure 5; Tables S5, S6). As an example, the ABC transporter *yadG* (AZ78_2378) was up-regulated by LeDSF3, IND, and GLY. Likewise, *lapE* gene (AZ78_3791), responsible for the secretion of the RTX adhesin LapA, was up-regulated by LeDSF3, IND, GLY, BUT and 4HBA. Type II secretion system (T2SS) genes, such as *gspG* (AZ78_4200) and *gspH* (AZ78_4201), were up-regulated by LeDSF3, IND, GLY, and BUT (Figure 5; Tables S5-S8). On the contrary, the expression of Type III secretion system (T3SS) was down-regulated by GLY and 3HBA (Figure 5; Tables S7, S9). Genes associated with type IV secretion system (T4SS), such as the duplicate gene pair located through the genome *virB6* (AZ78_0512) was down-regulated by LeDSF3, IND and GLY (Figure 5; Tables S5-S7). Additionally, IND repressed *virB5b* (AZ78_0703) and *virB3* (AZ78_0706), from the *virB1-virB11* complex, and the duplicates gene pairs located through the genome *virB5b* (AZ78_4957) and *virB6* (AZ78_0514 and AZ78_4958). ShlB from the two-partner secretion of Type V secretion system (T5bSS) was down-regulated by BUT and 4HBA (Figure 5; Tables S8, S10). Finally, DCSs regulated general secretory (Sec) pathways, such as *secB* (AZ78_2696) up-regulated by IND and BUT (Tables S6, S8), or *secF* (AZ78_4807) down-regulated by LeDSF3 and BUT (Figure 5; Tables S6, S8).

## Discussion

The behaviour of bacterial species mainly relies on communication systems [2] and many secreted metabolites characterising the cooperation among microorganisms, as well as antibiotics and toxins involved in microbial competition, are controlled by DCSs [48–50]. DCSs are not only involved in signalling among self-cells, but also in the detection of specific cues produced by other strains or species [48]. In fact, many bacterial species have receptors for DCSs that are not produced by the same species, such as LuxR *solos*, and abundant two-component signalling systems [48]. Genome mining results indicate that AZ78 and *Lysobacter* spp. may (at least) produce DSFs, IND, DFs and perceive DSFs, IND, DFs, and AHLs. As a consequence, DCSs (mainly IND and GLY) influenced AZ78 antibacterial and antioomycetes activity *in vitro*. Different intra-, interspecies and interkingdom DCSs might be used as cues for AZ78 to favour the regulation of molecular pathways related to cell persistence in the rhizosphere or for coercion (Figure 6). Thus, transcriptome profiles showed that DCSs might contribute to alert AZ78 against toxic compounds in the rhizosphere by triggering the expression of genes encoding efflux pumps that could actively extrude antibiotics, heavy metals, biocides, and solvents [51]. In addition, DCSs might help cells to escape from adverse conditions or to reach nutrients [52]. For example, LeDSF3, GLY, and BUT up-regulated the expression of genes related to the biogenesis of T4P involved in twitching motility. T4P-driven twitching motility is involved in a variety of physiological and social behaviours of a wide range of bacteria [53, 54]. For instance, twitching motility is required for colonization and infection of phytopathogenic fungi and oomycetes in *Lysobacter* spp. [55, 56] and it seems to be a DSF-dependent trait in *L. brunescens* and *L. enzymogenes* [9, 20, 21]. Interestingly, up-regulation of T4P by GLY and BUT came along with increased antimicrobial activity, suggesting that AZ78 might colonize new niches and form a stable community to outcompete other (micro)organisms upon the perception of these DCSs. Moreover, GLY and BUT down-regulated the transcription of tRNA genes. In *E. coli* down-regulation of tRNA was associated with the ability of this bacterium to cope with amino acid starvation (reviewed in [57]) and oxidative stress [58], which prompted us to think that GLY and BUT might enhance the stress response in AZ78 to promote survival and adaptation in the rhizosphere. In contrast, IND caused a dysregulation of T4P genes in AZ78 with possible losses of T4P functionality. Accordingly, IND decreases motility and biofilm formation in *E. coli* [59–62], probably as a manner to save energy and regulate growth dynamics [63]. In support to this hypothesis, IND diminished the AZ78 cell growth, although it was not possible to formulate a clear conclusion as knowledge about IND functions is contrasting [64–66].

**Figure 6.**
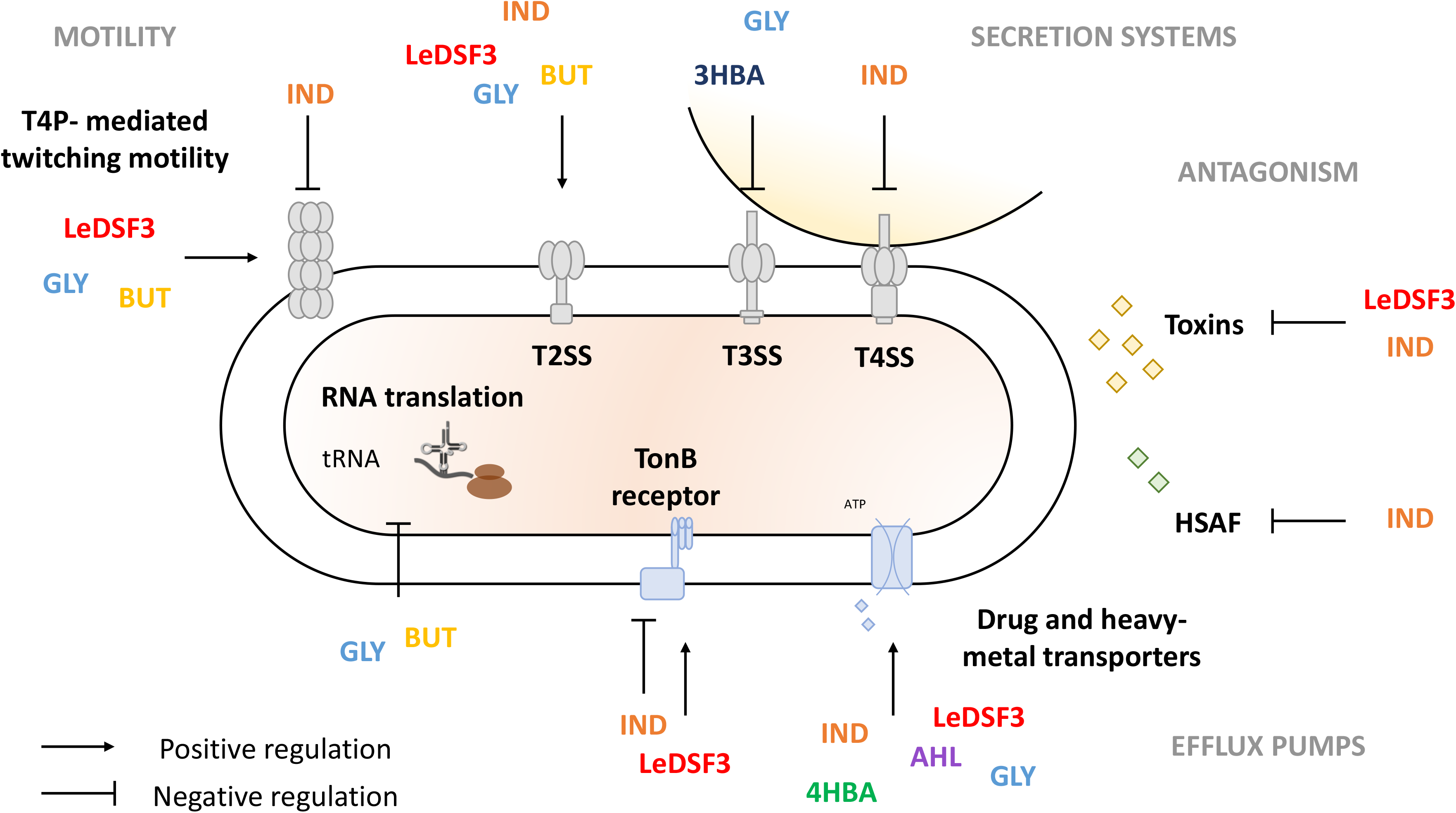
Schematic representation of *Lysobacter capsici* AZ78 response to diffusible communication signals. LeDSF3: 13-methyltetradecanoic acid; IND: indole; GLY: glyoxylic acid; BUT: 2,3-butanedione, 3HBA: 3-hydroxybenzoic acid; 4HBA: 4-hydroxybenzoic acid; AHL: N-acyl homoserine lactones.

In addition to escape mechanisms, *Lysobacter* cells might outcompete other microorganisms by producing extracellular lytic enzymes and antibiotics. Previous findings showed that HSAF biosynthesis is positively regulated by DSF, IND and 4HBA in *Lysobacter* spp. [9, 19, 21]. Yet, in AZ78 IND reduced antioomycete activity and down-regulated the expression of the HSAF biosynthetic gene cluster. The down-regulation of HSAF related genes by IND might be associated to the simultaneous up-regulation of the solo LuxR *solo* (AZ78_4823), as previously reported in *L. enzymogenes* OH11, where overexpression of *lesR* (LuxR homologue) leads to a decrease in HSAF production [67, 68]. Moreover, IND up-regulated several transcription regulators (among which various *tetR* repressors [69], like AZ78_0770 and AZ78 3232) that might be involved in HSAF biosynthesis regulation in AZ78, as found for LetR (a TetR-family protein) in *L. enzymogenes* OH11 [70]. The expression of HSAF biosynthetic cluster is also negatively regulated by cyclic-di-GMP (c-di-GMP) in *L. enzymogenes* OH11 [52]. Interestingly, IND up-regulated the expression of a diguanylate cyclase (AZ78_4062) that might be involved in c-di-GMP biosynthesis, implying a regulation role of c-di-GMP in HSAF production in AZ78. Besides producing secondary metabolites with antimicrobial activity, AZ78 might produce diffusible proteinaceous toxins and toxins deployed by contact-dependent systems, such as Rhs toxins, which mediate growth inhibition of neighbouring cells in *Dickeya dadantii* [71], or R-bodies, which are responsible for cell membrane disruption and toxins delivery in several bacterial genera [72, 73]. Thus, the down-regulation of several *rhs* and *reb* genes required for Rhs toxins and R-bodies synthesis by LeDSF3 and IND might have contributed to lower AZ78 the antioomycete and antibacterial activity. Moreover, AZ78 down-regulated signal transduction pathways in presence of IND, such as TonB-dependent receptors, which play a key role in microbial competition with the uptake of iron-siderophore complex or vitamins [74].

Bacteria often use secretion systems to manipulate and kill rival bacterial and eukaryotic cells [75, 76]. Of those, T3SS, T4SS and T6SS are related to the establishment of pathogenic interactions with microbial eukaryotic hosts in *Lysobacter* spp. [77]. Thus, modulation of genes related to secretion systems might result in gain/loss of ability to compete with other (micro)organisms [78]. In agreement with this statement, IND down-regulated T4SS and decreased antimicrobial activity in AZ78. Down-regulation of T4SS by IND might be related to the overexpression of diguanylate cyclases (e.g., AZ78_4062), responsible for c-di-GMP increase and T4SS inactivation in *A. tumefaciens* [79]. However, T3SS was down-regulated by GLY and 3HBA with no decrease in AZ78 toxic activity, suggesting that it was probably repressed to save energy under conditions where it does not provide an advantage, as found in *P. aeruginosa* [80]*, Vibrio harveyi* [81] and *Yersinia pseudotuberculosis* [82].

Overall, functional and transcriptome analysis of AZ78 in presence of signalling communication systems clarified its behaviour in the rhizosphere. Our results show that GLY and BUT might facilitate rhizosphere competence of *Lysobacter* spp. and soil disease suppressiveness to plant pathogens. On the other hand, IND might prevent *Lysobacter* spp. from growing at high cell densities and decrease motility. Moreover, IND and LeDSF3 might decrease *Lysobacter* spp. ability to control phytopathogenic microorganisms. Manipulating DCSs levels in the rhizosphere could therefore provide efficient means to favour the establishment and functioning of beneficial bacteria, such as *Lysobacter* strains, at the root-soil interface.

## Acknowledgements

This work was founded by the European Union’s Horizon 2020 research and innovation programme under the Marie Skłodowska-Curie grant agreements no. 797028.

## Author contributions

AB and GP conceived the study, performed the experiments, analysed the data, conceptualized, and drafted the manuscript. MP helped in the experimental set up and provided input and proof reading of the manuscript. IP provided input and proof reading of the manuscript.

