## Supplemental information for "Listening to bacterial Esperanto: transcriptome reprogramming in a plant beneficial rhizobacterium"

### Supplementary information

#### Supplementary figures

***rpfG***

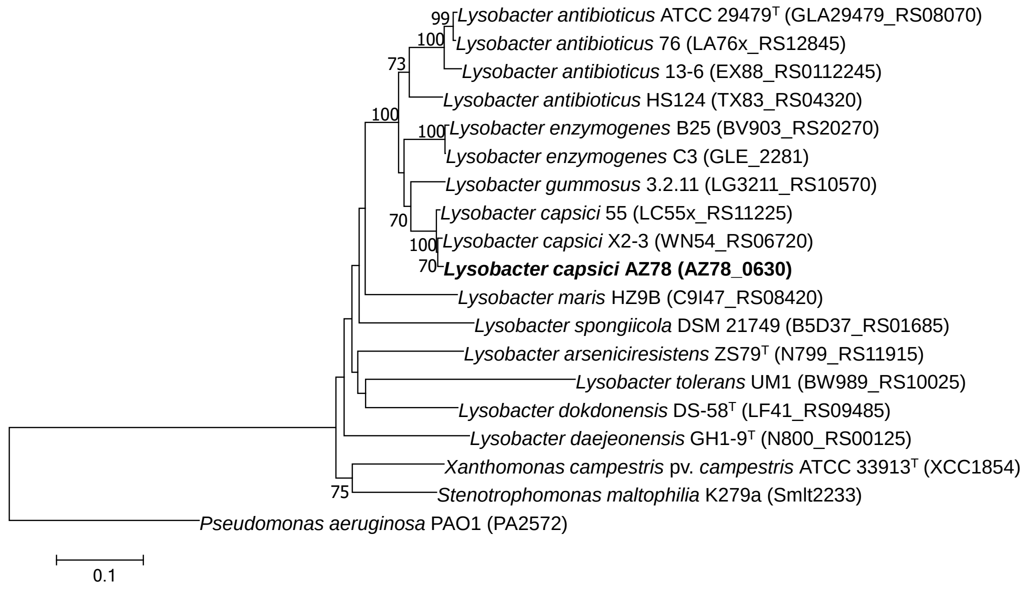


***rpfB***

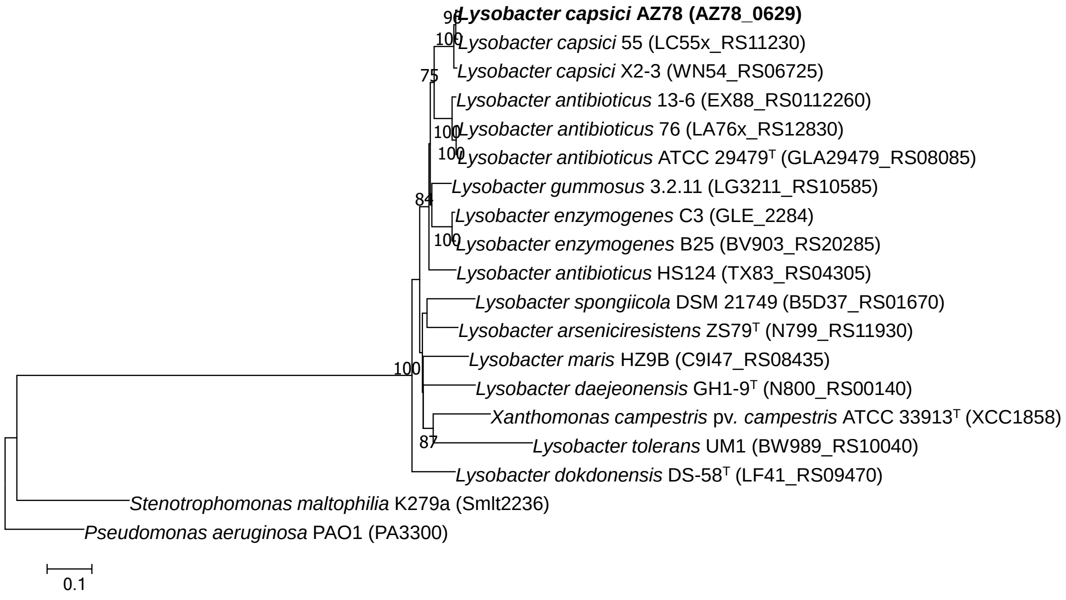


Figure S1. Neighbor-joining trees illustrating the relationships across *Lysobacter* members based on nucleotide sequences of the *rpfG* and *rpfB* genes. Locus tag numbers are given in brackets, GenBank accession numbers for the whole genome sequences are given in Table S1.

***rpfC***

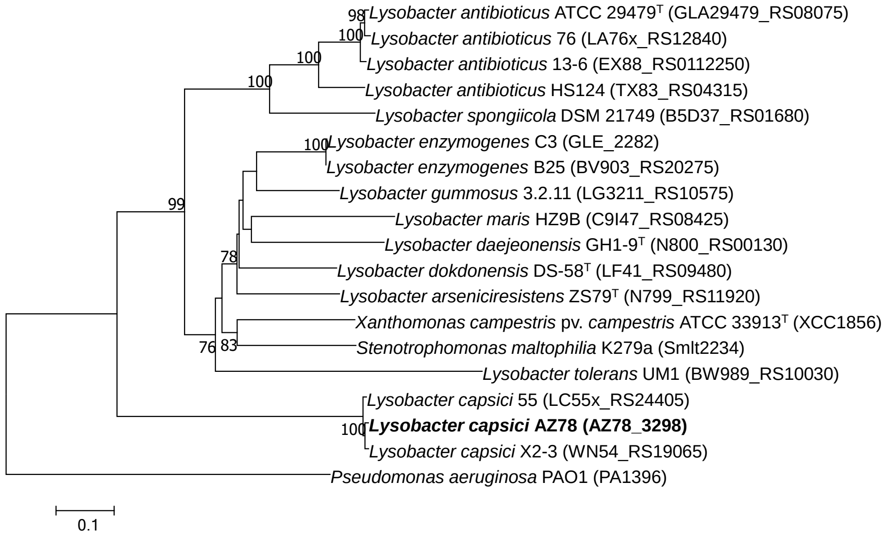


***rpfF***

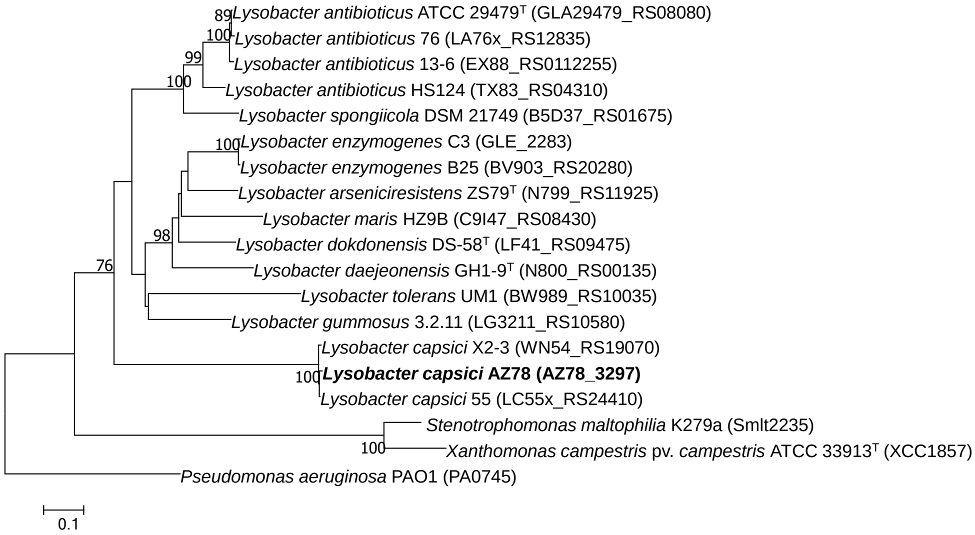


Figure S2. Neighbor-joining trees illustrating the relationships across *Lysobacter* members based on nucleotide sequences of the *rpfC* and *rpfF* genes. Locus tag numbers are given in brackets, GenBank accession numbers for the whole genome sequences are given in Table S1.

***trpC***

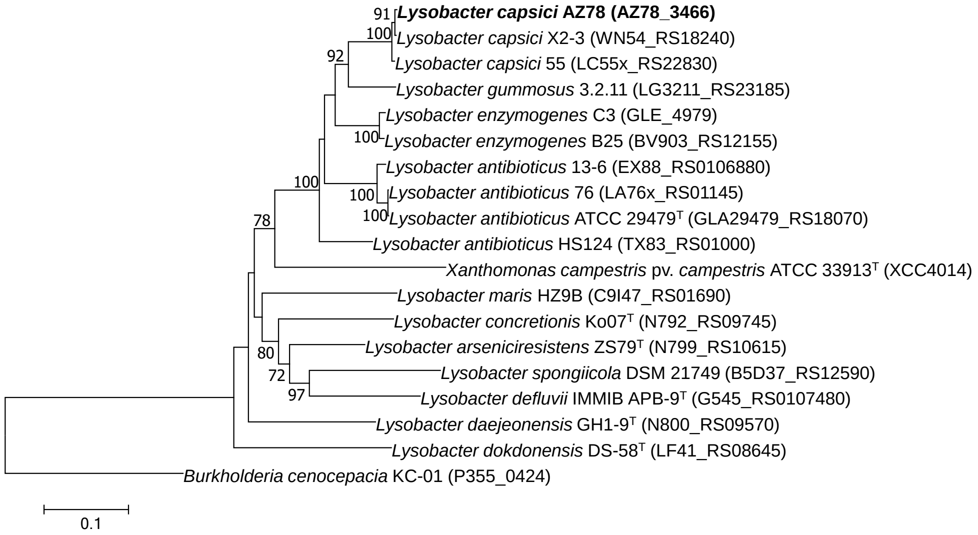


13

***qseB***

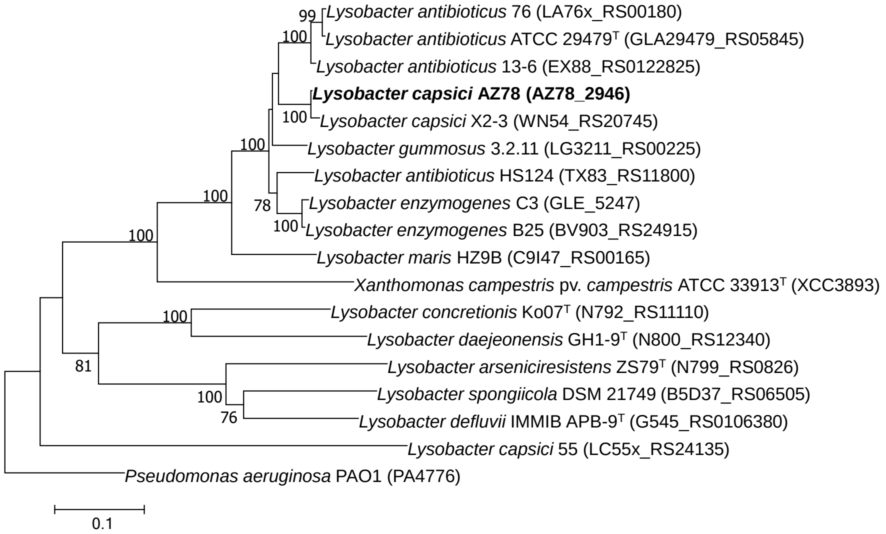


14

***qseC***

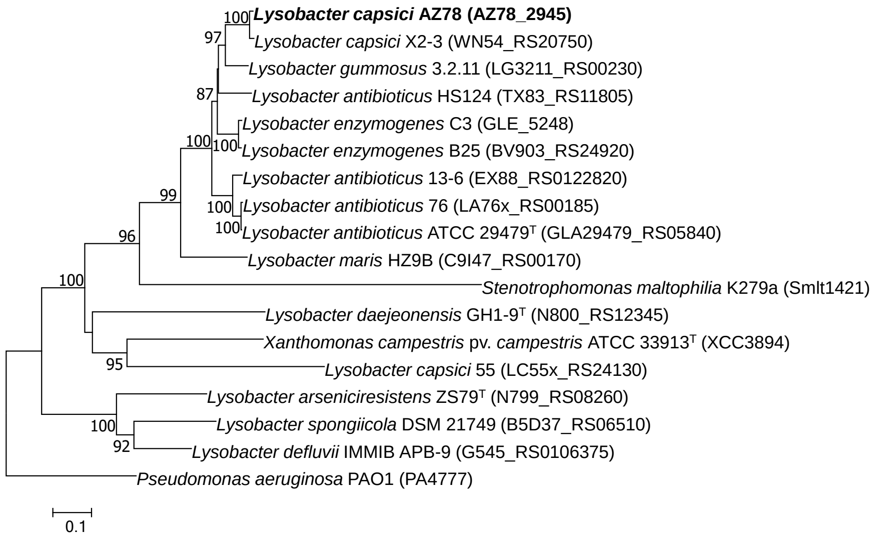


15

Figure S3. Neighbor-joining trees illustrating the relationships across *Lysobacter* members based on nucleotide sequences of the *tprC, qseB,* and *qseC* genes. Locus tag numbers are given in brackets, GenBank accession numbers for the whole genome sequences are given in Table S1.

***xanB2***

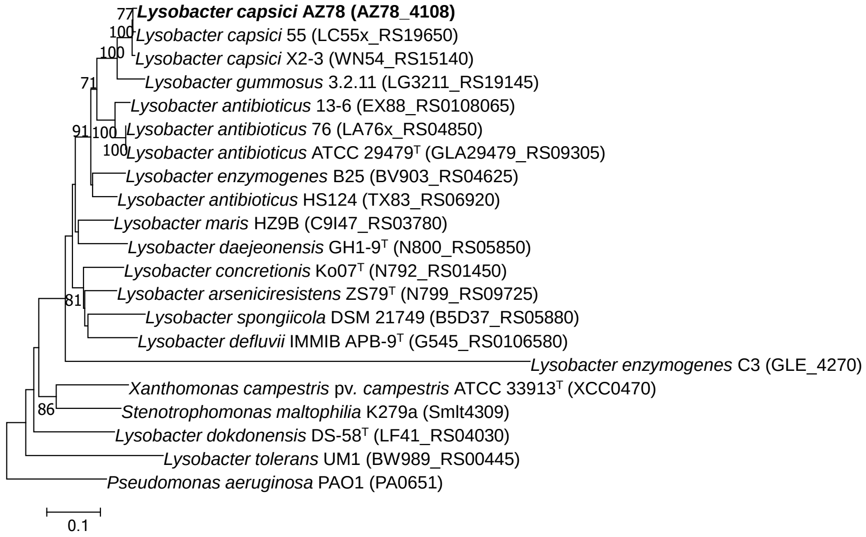


20

***lysR***

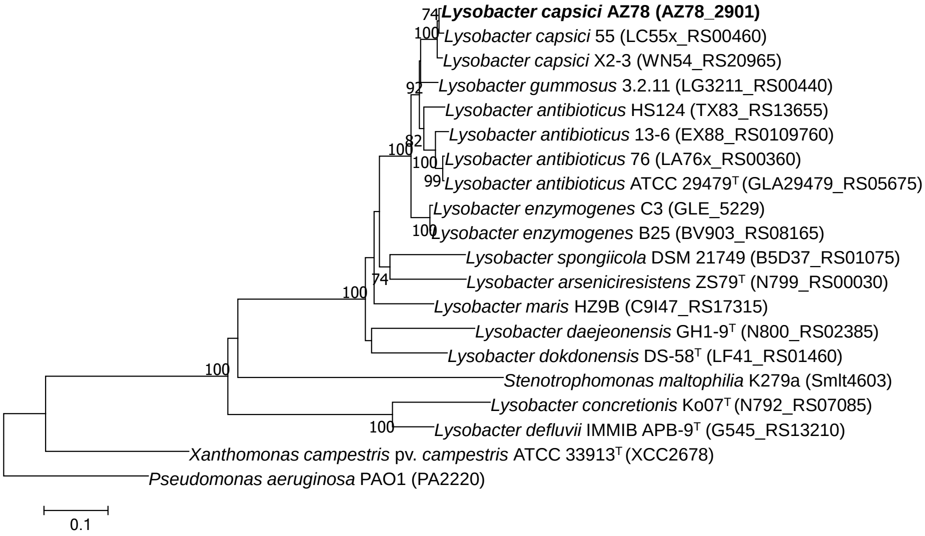


21

***luxR***

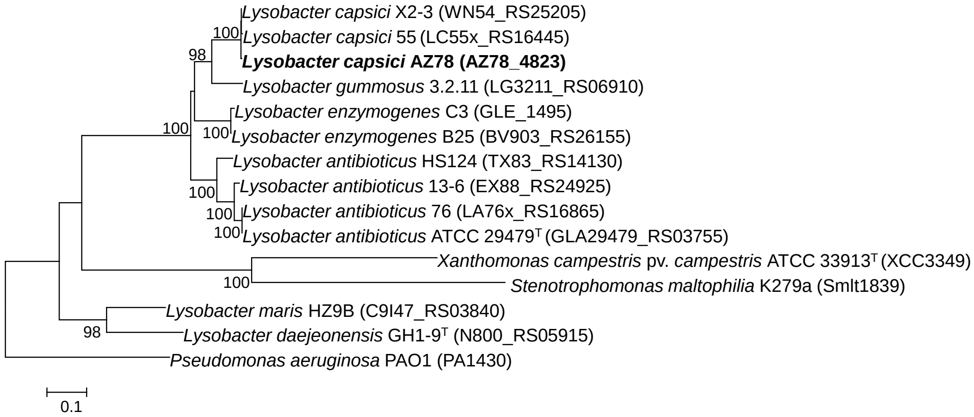


22

Figure S4. Neighbor-joining trees illustrating the relationships across *Lysobacter* members based on nucleotide sequences of the *xanB2, lysR,* and *luxR* genes. Locus tag numbers are given in brackets, GenBank accession numbers for the whole genome sequences are given in Table S1.

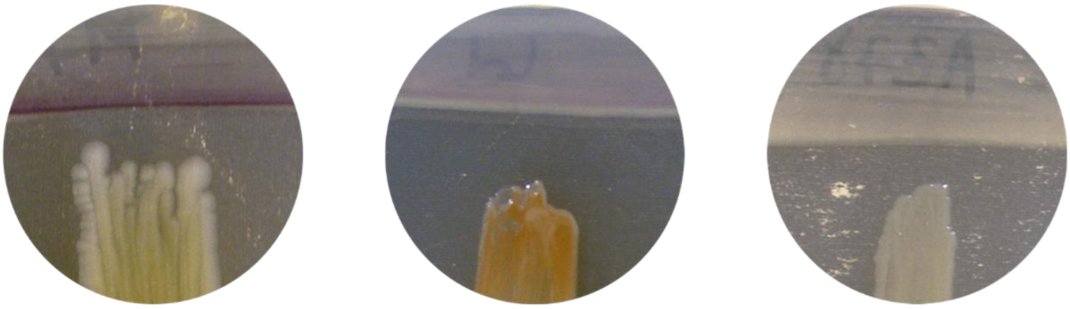

***Pseudomonas chlororaphis* M71** ***Lysobacter daejeonensis* GH1-9$^{T}$** ***Lysobacter capsici* AZ78**

Figure S5. Bioassay of N-acyl-homoserine lactones. N-acyl-homoserine lactones are produced by *Lysobacter daejeonensis* GH1-9$^{T}$ (purplish colour in the reporter strain *Chromobacterium violaceum* CV026), but not by *L. capsici* AZ78.

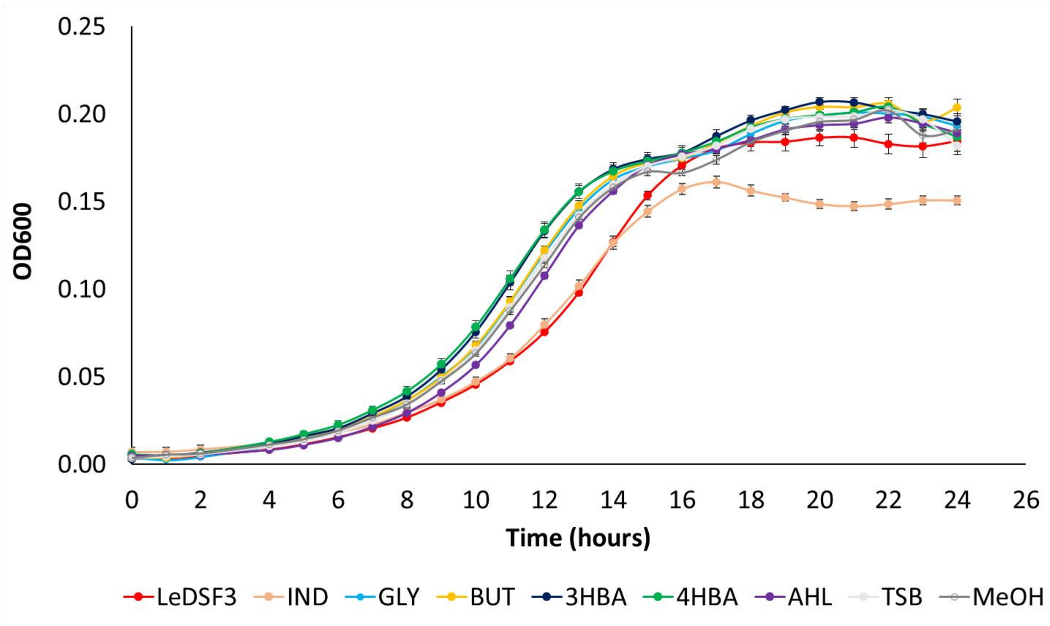


Figure S6. Growth curves of *Lysobacter capsici* AZ78 exposed to diffusible communication signals. LeDSF3: 13-methyltetradecanoic acid 50 µM, IND: indole 500 µM, GLY: glyoxylic acid 0.01 µM, BUT: 2,3-butanedione 0.01 µM, 4-HBA: 4-hydroxybenzoic acid 50 µM, 3-HBA: 3-hydroxybenzoic acid 30 µM, AHL: mix of N-acyl homoserine lactones 20 µM, TSB: 1/10 Tryptic Soy Broth, MeOH: 1% v/v methanol.

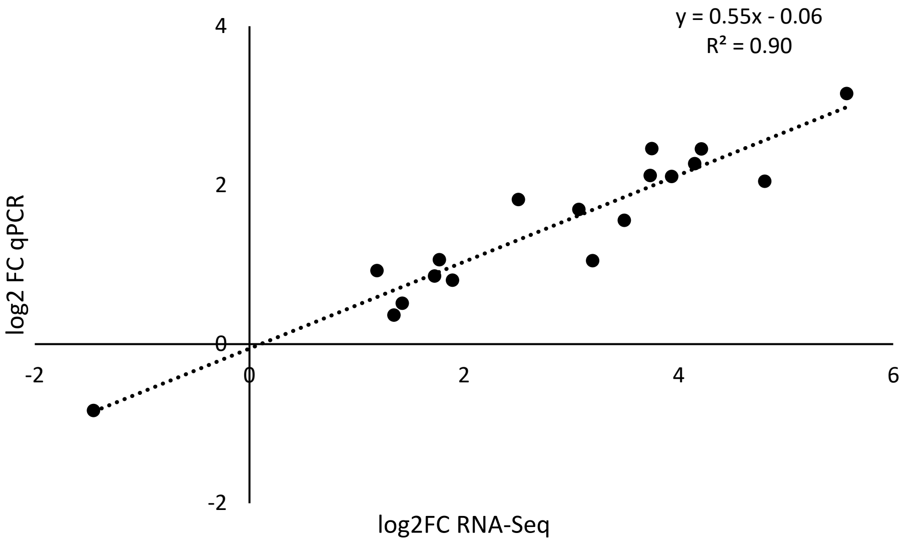


Figure S7. Scatter plot of RNA-Seq and qRT-PCR relative expression levels. Pearson correlation test (*r* = 0.95) was applied to log2 fold change (FC) values of selected genes (Table S4).

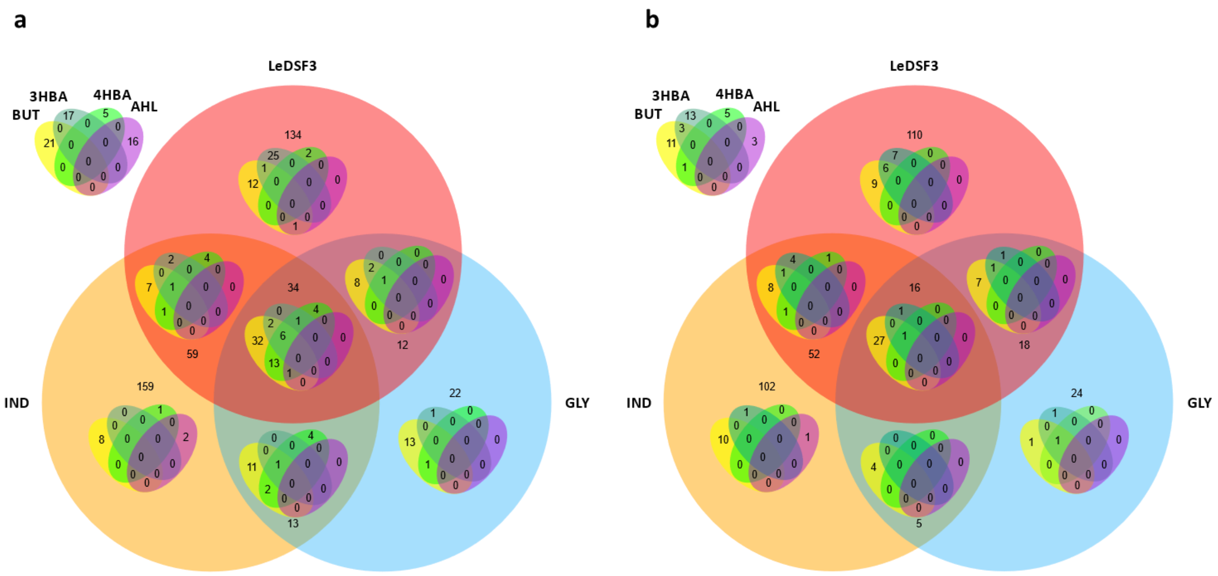


Figure S8. Venn diagram of up-regulated (a) and down-regulated (b) genes indicating the overlap in the number of differentially expressed genes (DEGs) in *Lysobacter capsici* AZ78 as response to diffusible communication signals. Only genes with |log2-fold change| > 1 and p-value < 0.01 were included. LeDSF3: 13-methyltetradecanoic acid 50 µM, IND: indole 500 µM, GLY: glyoxylic acid 0.01 µM, BUT: 2,3-butanedione 0.01 µM, 4-HBA: 4-hydroxybenzoic acid 50 µM, 3-HBA: 3-hydroxybenzoic acid 30 µM, AHL: mix of N-acyl homoserine lactones 20 µM.

#### Supplementary tables in the Excel file

Table S1. Bacterial strains used for phylogenetic analysis.

Table S2. Culture conditions of *L. capsici* AZ78.

Table S3. Number of reads obtained from Illumina HiSeq sequencing of RNA extracted from different *Lysobacter capsici* AZ78.

Table S4. Primers used in qRT-PCR.

Table S5. List of differentially expressed genes of *Lysobacter capsici* AZ78 after 48 h incubation with 13-methyltetradecanoic acid 50 µM.

Table S6. List of differentially expressed genes of *Lysobacter capsici* AZ78 after 48 h incubation with indole 0.5 mM.

Table S7. List of differentially expressed genes of *Lysobacter capsici* AZ78 after 48 h incubation with glyoxylic acid 0.01 µM.

Table S8. List of differentially expressed genes of *Lysobacter capsici* AZ78 after 48 h incubation with 2,3-butanedione 0.01.

Table S9. List of differentially expressed genes of *Lysobacter capsici* AZ78 after 48 h incubation with 3-hydroxybenzoic acid 30 µM.

Table S10. List of differentially expressed genes of *Lysobacter capsici* AZ78 after 48 h incubation with 4-hydroxybenzoic acid 50 µM.

Table S11. List of differentially expressed genes of *Lysobacter capsici* AZ78 after 48 h incubation with mix of N-acyl homoserine lactones 20 µM.
